# Ru(II)-Photoactive Agents for Targeting ER Stress and Immunogenic Cell Death

**DOI:** 10.1101/2024.07.18.604104

**Authors:** Madeline Denison, Alexander Ullrich, Mackenzie K. Herroon, Shane Mecca, Claudia Turro, Izabela Podgorski, Heather Gibson, Jeremy J. Kodanko

## Abstract

Immunotherapy has emerged as a promising avenue for cancer treatment by bolstering the immune system’s ability to recognize and attack cancer cells. Photodynamic therapy shows potential in enhancing antitumor immunity, though the mechanisms behind its success are not fully understood. In this manuscript, we investigate two previously reported green light activated PCT/PDT agents where compound **2** - [Ru(tpy)(Me2bpy)(**3**)]^2+^, (tpy = 2,2’:6’,2’’- terpyridine, Me2bpy = 6,6’-dimethyl-2,2’-bipyridine, **3** = pyridyl-BODIPY-I2,) - shows remarkable photoselectivity in assays containing both 2D cancer cells and 3D cocultures containing BALB/c macrophages and 4T1 murine breast cancer cells. Through flow cytometry and protein analysis, we found complex **2** displays superior evidence of induced endoplasmic reticulum (ER) stress markers and indicators of immunogenic cell death (ICD) compared to its ligand **3**, despite its weaker photoselectivity. Most importantly, these results were supported by *in vivo* studies where **2** produced anti-tumor immunity against the 4T1 tumor model in BALB/c mice. Complete tumor elimination was achieved in 2/8 mice, and these mice were both protected against a subsequent contralateral rechallenge and showed increased *ex vivo* peripheral tumor antigen-specific recall, suggesting memory T cells are induced by **2**. Signatures of M1 macrophage polarization were also evident in tumor tissue from the remaining 6/8 mice treated with **2** compared to untreated tumors.

These findings demonstrate Ru(II) complexation plays a critical role in ER targeting which triggers ICD, highlighting the potential of Ru(II) agents as future *in situ* tumor vaccines.

## Introduction

Cancer is the second leading cause of death worldwide, accounting for one in six deaths in 2020.^1^ Immunotherapy has evolved into an improved means for cancer treatment by enhancing the immune system’s recognition of cancer cells. By programming immune cells to attack cancer cells, this noninvasive treatment can offer successful, long-lasting remission, which is not achieved by common cancer treatment methods such as surgery, radiation therapy, and chemotherapy. Unfortunately, few patients initially respond to immunotherapy drugs, and those that do often form drug resistance.^2–4^ Alternatively, use of photodynamic therapy (PDT) has been shown to enhance systemic antitumor immunity, leading to tumor elimination and protection from recurrence.^5–8^ For instance, ruthenium (II) based PDT agent TLD-1433, currently in human clinical trials, was found to display antitumor immunity in a mouse model of colon cancer.^9^ However, factors that led to this response are still not understood. Additionally, our labs also found cotreatment of PCT/PDT agent [Ru(tpy)(Me2dppn)(py)](PF6)2 (**1**, Figure 1), where dppn = 3,6-dimethylnaphtho[2,3- *f*][1,10]phenanthroline, along with doxorubicin displayed synergistic signs of tumor-associated macrophage killing and immunogenic cell death (ICD), indicated by membrane translocation of calreticulin (CRT). While these results were promising, this light-activated approach involving **1** was achieved with blue light, which is known to exhibit minimal tissue penetration. In this manuscript, we report the detailed investigation of biologically relevant metalloimmunotherapy using agents activated with green light, and the consequential discovery of the importance of Ru(II) complexation for achieving ER-targeting and immunogenic cell death.

**Figure 1.**
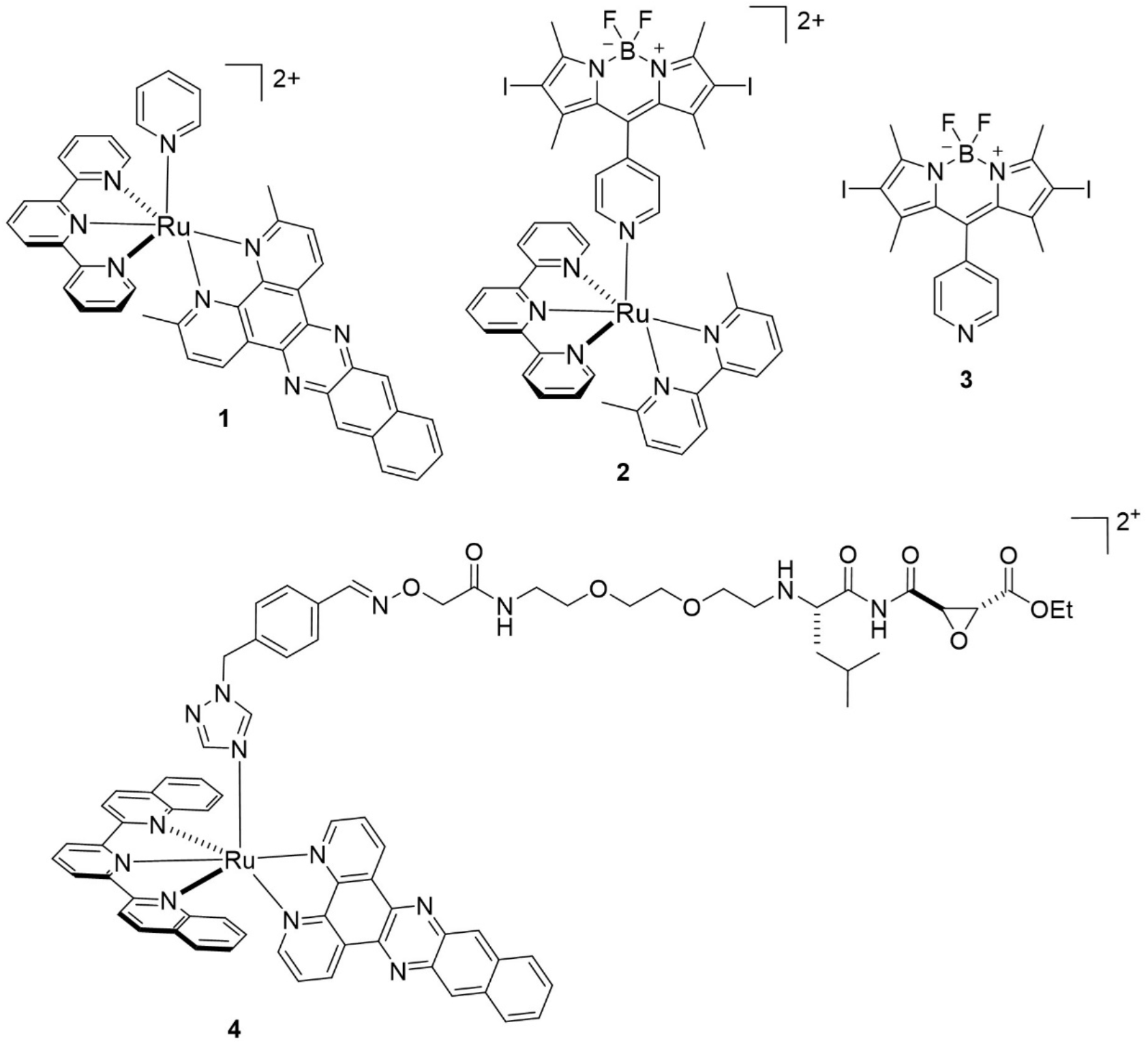
Previously reported Ru(II)-based PCT/PDT agents **1**, **2**, and **4** and BODIPY photosensitizer **3.**

Our initial studies began with two previously reported Ru(II)-based dual action PCT/PDT agents **2** and **4** (Figure 1).^10,11^ While both complexes are activated by green light and exhibit photoselective cell death in tumor cells, their photolabile ligands contribute to different photophysical or biological properties. Complex **2** incorporates a BODIPY (4,4-difluoro-4-bora-3a,4a-diaza-s-indacene) containing ligand **3** which attributes to a high molar extinction coefficient (ε542 = 52,900 M^−1^ cm^−1^) to allow for efficient light absorption. Conversely, complex **4** Ru(II) fragment [Ru(dqpy)(dppn)]^2+^, where dqpy = 2,6-di(quinolin-2-yl)pyridine and dppn = naphtho[2,3-*f*][1,10]phenanthroline, contributes to the absorption in the blue-green region, with a molar extinction coefficient ε470 = 9,800 M^−1^ cm^−1^. Complex **4** also contains a cysteine cathepsin inhibitor, which can serve as a delivery vector for the tumor microenvironment where cysteine cathepsins are overexpressed.^12^

## Results and Discussion

To begin our investigation, we first performed cellular viability experiments in murine breast cancer 4T1 cells. The 4T1 cell line was chosen to facilitate later translation to a syngeneic 4T1 BALB/c murine model where induction of adaptive anti-tumor immunity could be evaluated *in vivo*. The effects of compound **2** were determined in 4T1 cells plus or minus treatment with green light (520-25 nm, tirr =15 min) via the 3-(4,5-dimethylthiazol-2-yl)-2,5-diphenyltetrazolium bromide (MTT) assay (72 h). Cells were treated with various concentrations of **2** (0 – 500 nM) in quadruplicate wells and cells incubated for 4 h to allow for uptake, then washed and irradiated (520-25 nm, tirr = 15 min). After 72 h, viability was determined by MTT assay. Cellular viability was calculated for each concentration and data were fit to determine EC50 values (Figure 2). Experiments were run in triplicates to determine an average EC50 value. Complex **2** was found to have a light EC50 of 0.25 ± 0.02 μM, in contrast to 14.7 ± 0.8 μM (520-25 nm, tirr = 10 min) we reported previously for **4**.^11^ Importantly both complexes were found to be nontoxic at concentrations >50 μM in the dark, giving photoselectivity indices of >200 and >3 for complexes **2** and **4**.

**Figure 2.**
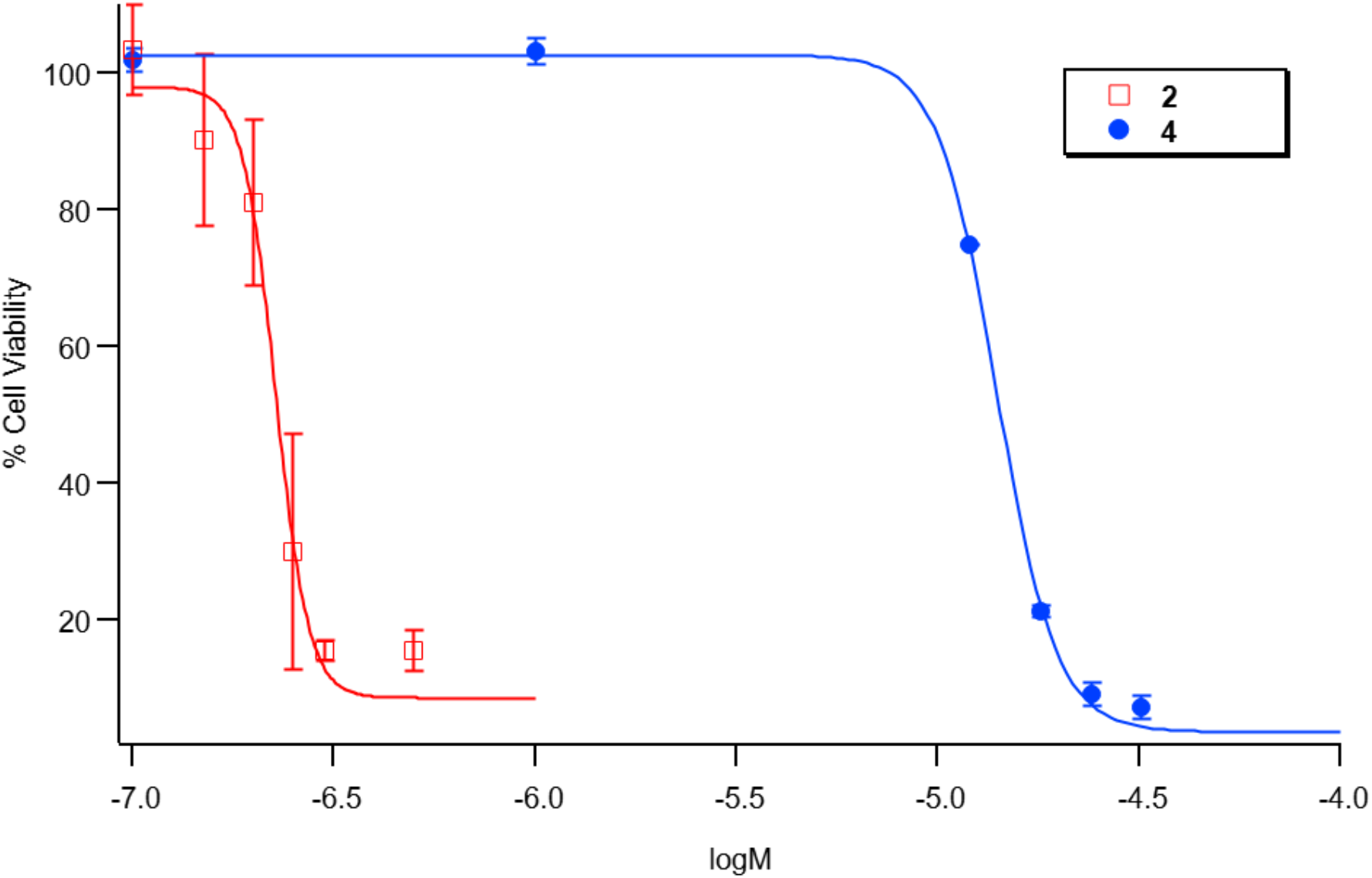
Representative EC50 plot depicting 4T1 cellular viability upon treatment of **2** and **4** under light conditions. Data for **4** were previously reported.^11^

Given promising photoselectivity in 2D 4T1 cells, we continued to study the efficacy of complexes **2** and **4** in 3D coculture containing macrophages and 4T1 tumor cells. Macrophages are the predominant immune cell in the tumor microenvironment, playing a crucial role in modulating the immune system through phagocytosis, antigen presentation, and the secretion of cytokines, chemokines, and growth factors. This diverse coculture offers a more precise representation of the tumor microenvironment, encompassing interactions with the extracellular matrix (ECM), cell polarity, and cell-to-cell contacts. As a preliminary screen, we first assessed viability of cells within the coculture following treatment of **2** and **4** by Alamar blue. After establishing the 3D coculture spheroids of 4T1 cells and BALB/c macrophages, spheroids were treated with various concentrations of **2** and **4** and incubated for 24 h to allow for cellular uptake. Media was changed and cells were then irradiated (520 – 25 nm, tirr = 15 min for **2**, tirr = 10 min for **4**). After 48 h, viability was assessed by Alamar blue. Unfortunately, we found complex **4** showed no effects on cell viability at ≤ 30 μM upon irradiation (data not shown). Solubility issues limited our ability to evaluate **4** at higher concentrations. These results were consistent with other Ru(II) agents that needed to be dosed at concentrations, as high as 70 to 90 μM, to elicit cell killing in 3D tumor spheroids.^13,14^ Fortunately, complex **2** showed a response at concentrations < 10 μM. We hypothesized this success was due to the red shifted absorption and nearly ten-fold larger extinction coefficient of complex **2** vs. complex **4** at 520 nm, where improved absorptivity leads to efficient PDT action. Because metabolic stress can lead to an inaccurate representation of cellular viability using metabolic assays such as MTT and Alamar blue,^15–17^ we chose to employ the live/dead assay which relies on cell membrane permeability. Calcein acetoxymethyl (AM) labels live cells upon hydrolysis by intracellular esterases, whereas cell impermeable ethidium homodimer-1 labels dead cells by binding to DNA. Using similar conditions to the Alamar blue assay, viability of tumor spheroids was assessed 48 h after irradiation treatment. Confocal microscopy was used to image live, green cells and red, dead cells followed by quantification using Volocity software. Remarkably, complex **2** showed significant cell killing in the 4T1/macrophage cocultured spheroids at concentrations as low as 1 μM under light conditions (Figure 3).

**Figure 3.**
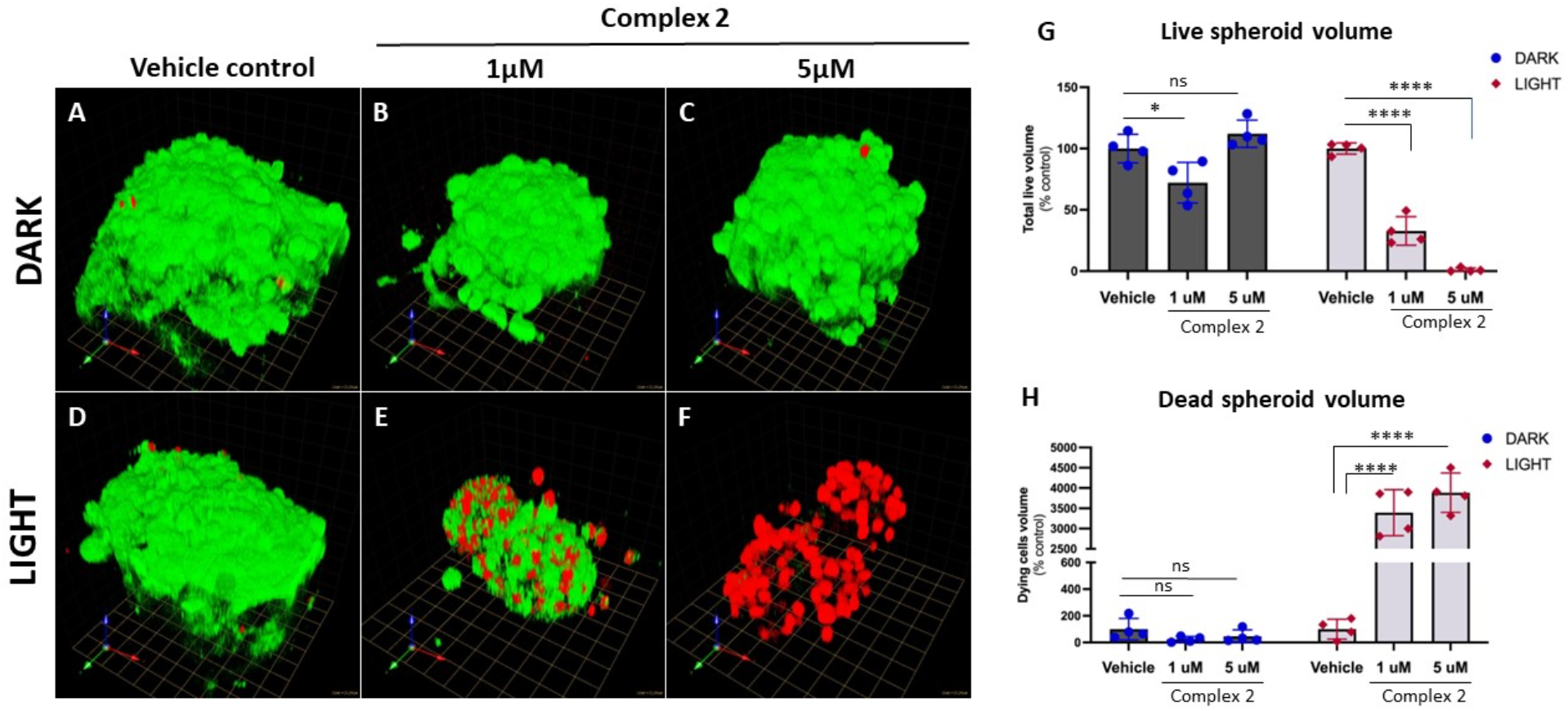
3D coculture spheroids stained with Calcein AM (green, live) and Ethidium Homodimer-1 (red, dead) were imaged on a Zeiss 780 Confocal microscope followed by volume quantification. Spheroids were treated with increasing concentrations of complex **2** left in the dark (**A-C**) or irradiated (**D-F**). Using Volocity, the live volume of the spheroids (calcein-positive) was calculated (**G**), and the percentage of dead cells (ethidium homodimer-positive) compared to the control dark conditions were calculated (**H**). **A-F** – 1 unit (1 box) = 21.34μm

We then assessed the potential of complex **2** to elicit ER stress by probing for damage-associated molecular patterns (DAMPs). Release of DAMPs, such as CRT, enhances the immunogenicity of the dying cells.^18,19^ In particular, the translocation of CRT to the cell surface during ER stress-induced ICD facilitates the recognition and engulfment of tumor cells by macrophages and dendritic cells (DCs), leading to T-cell-mediated elimination of the tumor.^20^ 4T1 cells were treated with varying concentrations of **2** (0-1 μM), incubated for 4 h, irradiated with green light (λirr = 520-25 nm; tirr = 15 min) or left in the dark, and incubated for an additional 4 h. Flow cytometric analysis of 4T1 cells revealed surface CRT exposure at ≥500 nM **2** under light conditions (Figure 4). Importantly, minimal CRT release was found in the dark indicating this stress response is light dependent.

**Figure 4.**
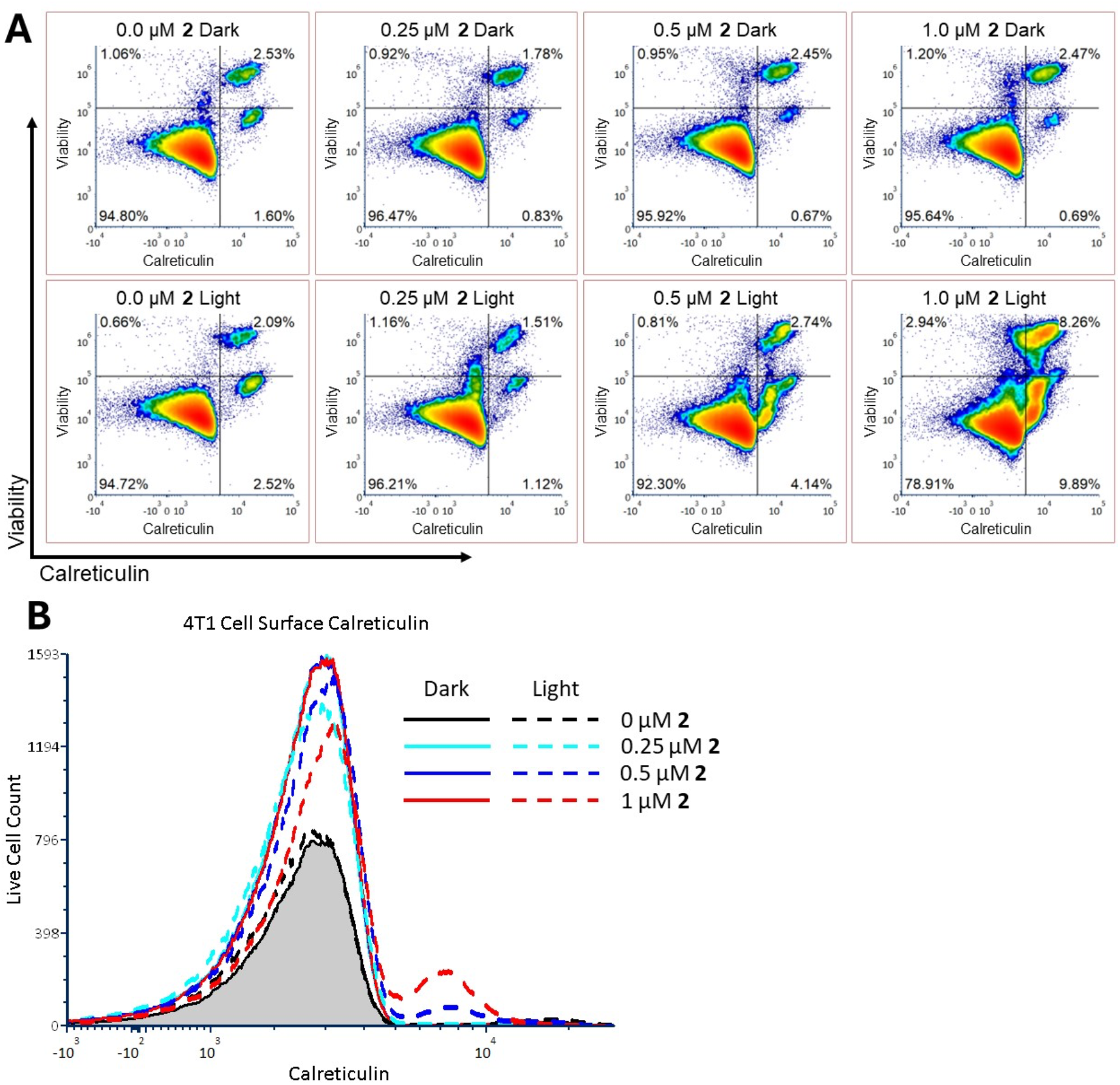
Mouse 4T1 cell calreticulin (CRT) cell surface exposure after *in vitro* treatment with compound **2** with and without light irradiation. Tumor cells were treated with increasing doses of **2** as indicated for 4 hours prior to green light irradiation. Cells were collected 4 hours later and stained with ViaDye Red viability dye and PE-conjugated anti-CRT with analysis by flow cytometry. **A)** Density plots of viability dye uptake versus CRT. **B)** Histograms of surface CRT on viable cells.

To further substantiate the role of **2** in ER stress, we investigated key proteins found in the unfolded protein response (UPR) signaling pathway. Under ER stress, the UPR is triggered to protect the ER against stress and reestablish homeostasis. Of the three arms that UPR is divided into, the protein kinase R-like endoplasmic reticulum kinase (PERK) arm is found to be strongly related to ICD and release of CRT.^21,22^ Thus, we chose to examine levels of transcription factors found in the PERK pathway, activating transcription factor 4 (ATF4) and C/EBP homologous protein (CHOP), in pretreated 4T1 cell lysates. 4T1 cells were treated with varying concentrations of **2** (0-500 nM), incubated for 4 h, washed, and irradiated with green light (λirr = 520-25 nm; tirr = 15 min) or left in the dark, and incubated for an additional 4 h before lysing. Protein analysis of lysates by western blot indicated ATF4 and CHOP were detected at concentrations as low as 100 nM under light conditions (Figure 5). Because structure-activity relationships of Ru and related metal complexes have been shown to facilitate ER targeting,^23–25^ we hypothesized the same levels of ER stress markers would not be apparent upon treatment with ligand **3**. To confirm this, 4T1 cells were concurrently treated with compound **3** and lysates were collected for western blot analysis. Minimal expression of ATF4 and CHOP were found in cells treated with **3** over the same concentration range. Critically, these results are consistent with the Ru(II) complex in **2** being crucial for driving ER stress and inducing ICD.

**Figure 5.**
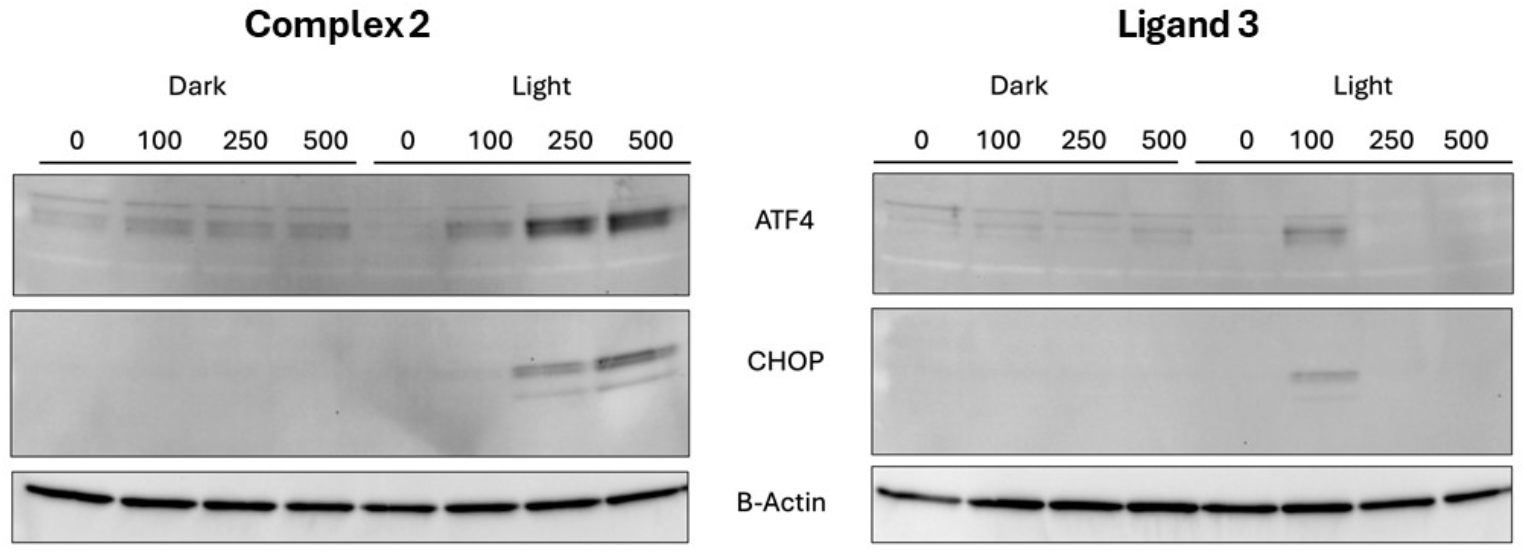
ATF4 and CHOP expression in murine 4T1 cells treated with varying concentrations (0 – 500 nM) **2**, **3**, or vehicle (0.1% DMSO) under dark or light conditions.

While studies successfully indicated complex **2** produces ER stress in 4T1 murine breast cancer cells, we sought to evaluate whether the same results could be found across human cancer cell lines. ER stress marker ATF4 was probed in MDA-MB-231 human triple negative breast cancer cells, and ATF4 and CHOP in DU-145 human prostate cancer cells. Cells were treated with **2** (0-500 nM) analogously to 4T1 cells, lysed, and analyzed by western blot. Again, ATF4 was evident in both light-treated MDA-MB-231 and DU-145 cells, with a strong response witnessed for CHOP in DU-145 cells (Figure 6). Collectively, these results verify ER stress following treatment of **2** is not restricted solely to murine breast cancer cells but can be found in human cells of various cancer types.

**Figure 6.**
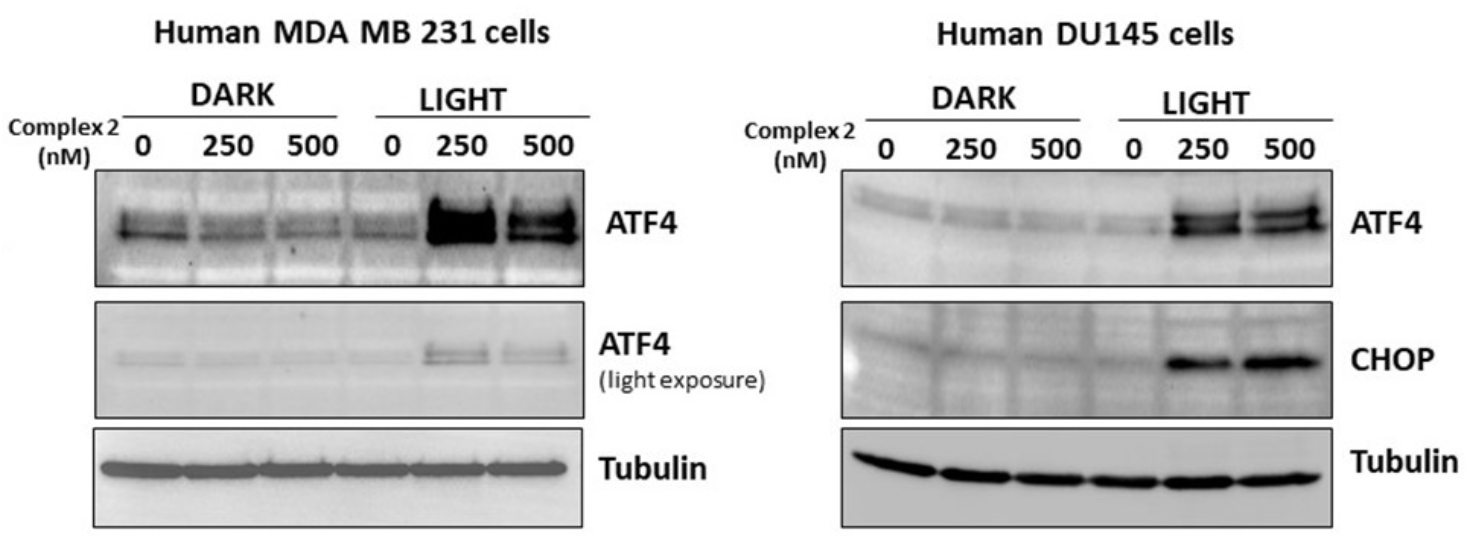
ATF4 and CHOP expression in human MDA-MB-231 and DU-145 cells treated with varying concentrations (0 – 500 nM) **2** or vehicle (0.1% DMSO) under dark and light conditions.

Finally, we assessed the capability of **2** to cause tumor ablation and induction of anti-tumor immunity *in vivo*. BALB/c mice were orthotopically inoculated with 4T1 cells stably expressing luciferase (4T1/Luc2), and when the tumor burden reached approximately 20 mm³, 10 μL of 0.5 mg/kg **2** (DMSO) was injected intratumorally. Four hours later, the tumor was pulsed twice for 10 minutes with a 532 nm laser, with a 10 minute delay between laser pulses. The treatment was administered for 3 cycles, given every 4th day, which resulted in complete tumor elimination in 2/8 treated mice (Figure 7A). To test for systemic adaptive antitumor immune memory, complete responder mice received a contralateral tumor rechallenge without further therapeutic intervention. Both mice were protected from the secondary tumor. After the rechallenge, spleens were collected for *ex vivo* immune analysis by interferon gamma ELISpot using irradiated 4T1/Luc2 cells as a source of tumor antigens. Splenocytes from the 2 complete responders had significantly increased tumor antigen recall relative to both untreated tumor-bearing mice and naïve mice, further indicating that *in vivo* treatment with **2** induced peripheral antitumor immune memory (Figure 8). Immunohistochemical (IHC) analysis of tumor outgrowth in the 6 non-responding mice as well as untreated control mice demonstrated no difference in macrophage (Mφ) infiltration (Figure 7B- D), however, RT-qPCR analysis of homogenized tumors revealed significantly upregulated IL-12 transcripts, indicating increased anti-tumor (M1) Mφ activity (Figure 7E).

**Figure 7.**
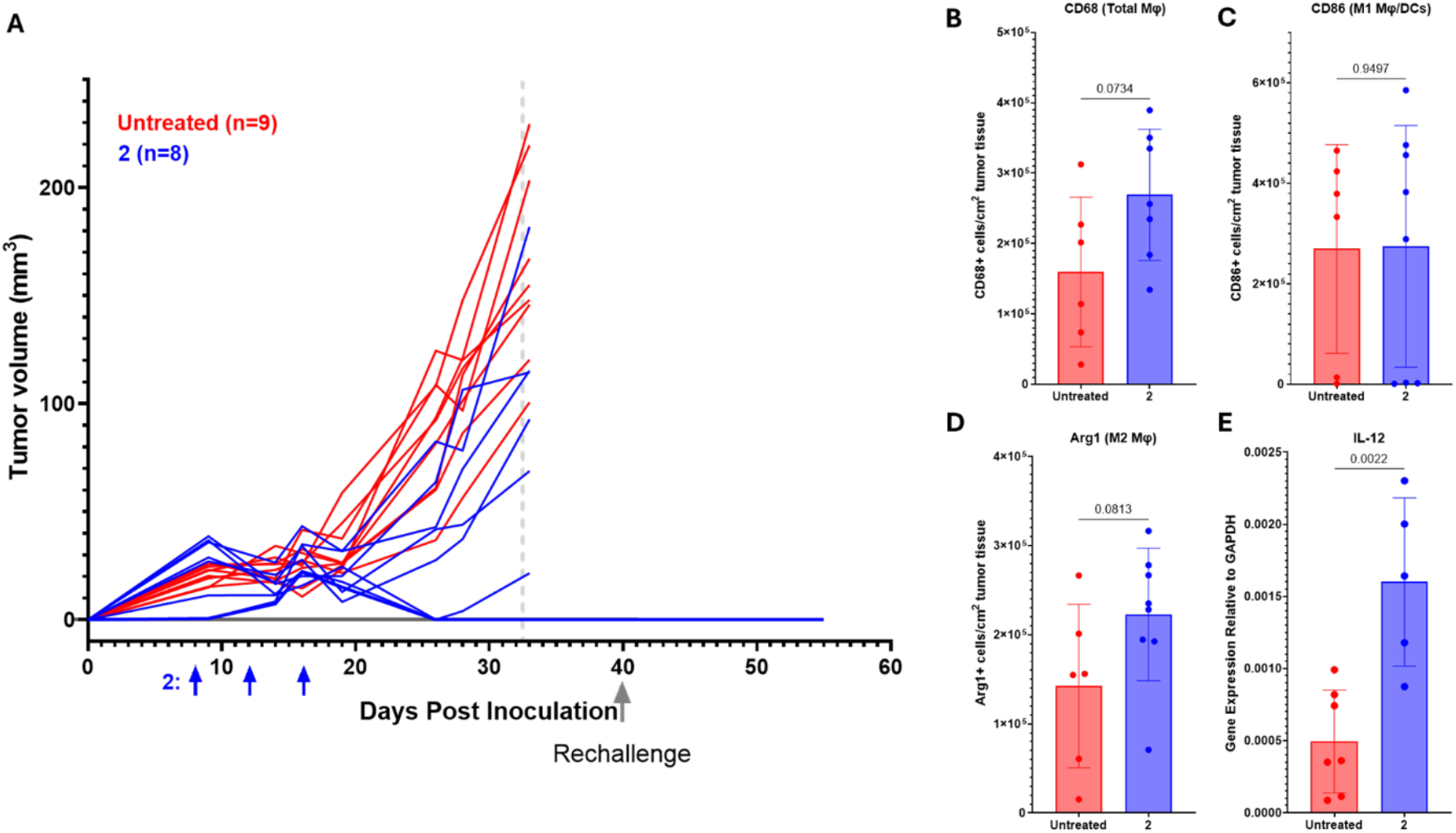
A) BALB/c mice were inoculated with 2x10^5^ 4T1/Luc2 cells orthotopically in the #4 mammary fat pad. Upon development of a palpable tumor on day 8, **2** was administered as 0.5 mg/kg every 4^th^ day for a total of 3 doses (blue arrows). Ten days after the last dose, a contralateral 4T1/Luc2 re-challenge was given without additional treatment. Untreated and non-responder treated mice were euthanized and tissues were collected 17 days after the last treatment. Responder mice were euthanized and tissues were collected two weeks after the rechallenge. IHC enumeration of **B)** CD68 (Total Mφ), **C)** CD86 (M1 Mφ and DCs), and **D)** Arg1 (M2 Mφ). **E)** RT-qPCR analysis of IL-12 expression.

**Figure 8.**
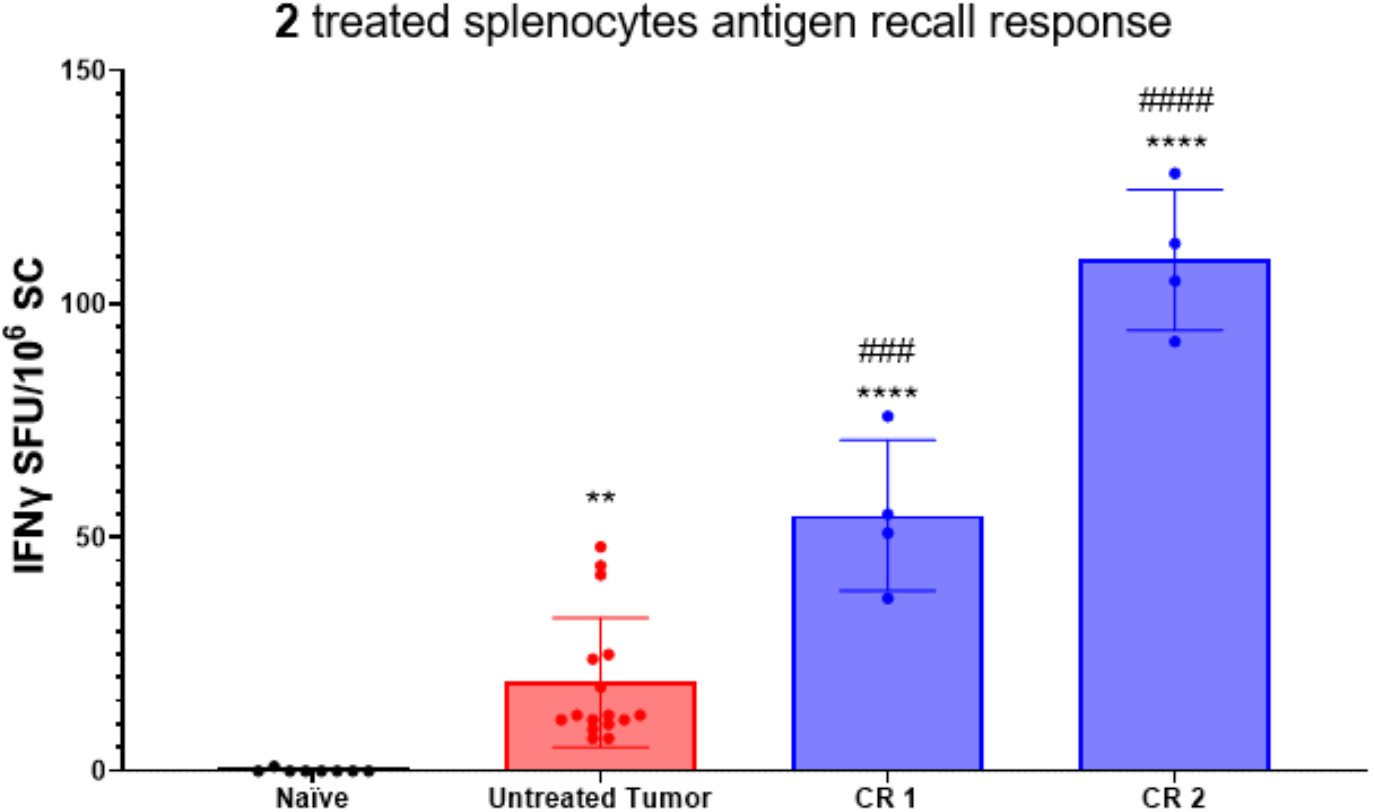
Splenocytes (SC) were collected from the 2 complete responder mice, as well as 4 untreated tumor-bearing mice and cultured with irradiated 4T1/Luc2 cells. IFNγ secreting SC were enumerated by ELISpot. Naïve BALB/c SC serve as controls.

In summary, we have shown complex **2** shows superior photoselectivity in 2D cancer cells and 3D coculture containing 4T1 and macrophage cells, likely due to its improved light absorptivity. While our previously reported complex **1** was successful in eliciting ICD markers such as CRT in combination with doxorubicin, complex **2** achieves analogous effects without an additional therapeutic agent. Importantly, this immunotherapeutic effect is obtained with green light which allows for better tissue penetration as opposed to blue light required in photoactivation of **1**. Additionally, our results indicate that complexation of ligand **3** to Ru(II), as in **2**, enhances ER stress, which may indicate a broader role for Ru(II) complexes as delivery vectors for strong absorbing dyes towards the ER to maximize ICD. Finally, we provide evidence that **2** is able to induce peripheral antitumor immune memory *in vivo*, protecting against tumor rechallenge.

Further investigations are ongoing that will evaluate the role of **2** and related compounds as *in situ* antitumor vaccines.

## Acknowledgments

We gratefully acknowledge the National Science Foundation (CHE 2102508), National Institutes of Health (T32GM142519), and Wayne State University (Grants Boost and Rumle Fellowship to MD) for support of this research.

## Experimental Procedures

### General Considerations

4T1 cells were maintained using RPMI 1640 medium containing 10% FBS, 100 units/mL penicillin-streptomycin (P/S), and 8 µg/mL blasticidin (InvivoGen). DU-145 and MDA-MB-231 cells were maintained using Dulbecco’s Modified Eagle Medium (DMEM) containing 10% FBS and 100 units/mL P/S. Macrophages from BALB/c mice were grown in a mix of the above RPMI media with 30% L929-conditioned media added as a source of M-CSF to allow the macrophages to grow. In the experimental conditions, the percentage of L929- conditioned media was reduced to 15%. All cells were kept in a humidified, 37°C, 5% CO2 incubator. Cells under light conditions were treated with green light (520-525 nm) for 15 mins at 25°C, whereas cells under dark conditions were left in the dark for 15 mins at 25°C.

### MTT Assay

4T1 cells were seeded at 7,000 cells per well in 100 µL growth media on a 96 well plate (Cell Treat) and allowed to incubate for 18 h in a 37°C humidified incubator ventilated with 5% CO2. Next, growth medium was aspirated and replaced with medium containing varying concentrations of **2** (0.05 – 50 µM) or vehicle (1% DMSO) in quadruplicates. Following a 4 h incubation period, medium was replaced with fresh medium, and cells were treated with light or left in the dark.

After incubating the cells for an additional 72 h, 10 µL of (3-(4,5-Dimethylthiazol-2-yl)- 2,5-Diphenyltetrazolium Bromide) (MTT, 5 mg/mL) reagent was added to every well. Cells were then allowed to incubate for 2 h. After this period, medium was aspirated, and resultant purple formazan crystals were dissolved in 100 µL DMSO. Plates were shaken for 20 min to ensure dissolution and absorbance was measured at 570 nm in every well. Absorbance of blank wells containing only DMSO were subtracted from the remaining vehicle and experimental wells. Cell viability at each concentration was calculated by dividing the corrected mean absorbance in wells containing compound by the mean absorbance of wells containing vehicle and expressing the ratio as a percentage value. For each experiment, cell viability vs. concentration was plotted and fitted to a sigmoidal curve to derive the cell viability at 50% or the EC50 value. Reported EC50 values are an average of triplicate experiments and error bars are shown as standard deviations.

### Live/dead Assay

3D spheroids were established using non-diluted reconstituted basement membrane (rBM; Cultrex™; R&D Systems) on acid-washed glass coverslips. An equal number of cells from both 4T1 murine cancer cells and macrophages from BALB/c mice (4,000 per cell line) were plated and spheroids were allowed to form. After 72 h, various concentrations of compound **2** were added to the media for 24 h. The spheroids were then carefully rinsed with PBS and had fresh media added before cells were treated with light or dark conditions. After 48 h, spheroids were stained with 2 μM Calcein AM and 5 μM Ethidium Homodimer-1 for 30 min at 25°C, washed with PBS, and immediately imaged by capturing z-stacks through the depth of the structure using a Zeiss LSM 780 Confocal Microscope with a 40× water immersion objective. Live cells (green; Calcein AM) were captured using excitation at 488nm and emission at 507nm. Dead cells (red; Ethidium Homodimer-1) were recorded using excitation at 488nm and emission at 730nm. 3D reconstruction and volume measurement per channel were quantified using Volocity Software (PerkinElmer).

### Flow Cytometry

Cells collected for flow cytometry were plated in 6-well plates the day prior to treatment. 4T1 cells were plated at 450,000/well in complete growth medium (2mL). Cells were allowed to incubate for 18 h in a 37°C humidified incubator ventilated with 5% CO2. Then, growth media was aspirated and replaced with medium containing various concentrations of **2** (0 – 1 μM). Following a 4 h incubation period, medium was replaced with fresh medium, and cells were irradiated or left in the dark. After incubating the cells for an additional 4 h, the cells were removed from the culture plates with trypsin and washed with PBS. The cells were then stained with ViaDye^TM^ Red viability stain (1:10,000) (Cytek Biosciences) and PE conjugated anti-Calreticulin (1:200) (Cell Signaling Technologies) prior to analysis on a Cytek Northern Lights^TM^ spectral cytometer in the Wayne State University Microscopy, Imaging, and Cytometry Resources (MICR) Core. FCS Express^TM^ 7 (De Novo Software) was used to analyze the results.

### Protein expression of ER stress markers

Cells collected for lysates were plated in 6-well plates the day prior to treatment. 4T1 cells were plated at 500,000/well, MDA-MB-231 and DU145 cells were plated at 1,000,000/well in their respective media (2 mL). Cells were allowed to incubate for 18 h in a 37°C humidified incubator ventilated with 5% CO2. Then, growth media was aspirated and replaced with medium containing various concentrations of **2** (0 – 500 nM), **3** (0 – 500 nM) or vehicle (0.1% DMSO). Following a 4 h incubation period, medium was replaced with fresh medium, and cells were irradiated or left in the dark. After incubating the cells for an additional 4 h, cells were washed three times with warmed PBS and lysed with 100 µL buffer containing 250 mM sucrose, 25 mM MOPS, 1 mM EDTA, 0.1% Triton X-100, and Halt™ Protease and Phosphatase Inhibitor Cocktail and frozen. Protein was analyzed by Bradford assay and Western blotting.

### In vivo experiments

BALB/c mice were inoculated with 2x10^5^ 4T1/Luc2 cells orthotopically in the #4 mammary fat pad. Upon development of a palpable tumor on day 8, **2** was administered as 0.5 mg/kg, followed 4 hours later with two 10 minutes with a 532 nm laser, with a 10 minute delay between laser pulses, every 4^th^ day for a total of 3 doses. Ten days after the last dose, 2x10^5^ 4T1/Luc2 cells were inoculated in the contralateral mammary fat pad of responder mice. This contralateral tumor re-challenge was given without additional treatment. Untreated and non-responder treated mice were euthanized and tissues were collected 17 days after the last treatment. Responder mice were euthanized, and tissues were collected two weeks after the rechallenge. The collected tissues were fixed in formalin and embedded in paraffin (FFPE) for IHC and snap frozen in liquid nitrogen and stored at −80°C for RT-qPCR.

### Immunohistochemistry

Sectioned FFPE tumor tissues were deparaffinized with HistoClear^TM^ (National Diagnostics) and rehydrated in an ethanol gradient prior to antigen retrieval. Antigen retrieval was performed with a citrate buffer (CD68 and Arg1) or a tris buffer (CD86). Tissue sections were blocked with BLOXALL^®^ (Vector Laboratories) for 30 minutes followed by normal horse serum for 1 hour prior to staining with CD68 (1:150) (Abcam), CD86 (1:800) (Invitrogen), or Arg1 (1:1000) (Proteintech) overnight at 4°C in a humidified chamber. The slides were then washed 2X with 1X PBS and alkaline phosphatase (AP) conjugated horse-anti-rabbit secondary antibody (Vector Laboratories) was incubated for 30 minutes at room temperature. Following another PBS wash, ImmPACT^®^ Vector^®^ Red AP Substrate (Vector Laboratories) was incubated on the tissues for either 5 minutes (CD68) or 10 minutes (CD86 and Arg1). Slides were then scanned and quantified with FIJI/ImageJ, utilizing the Trainable WEKA Segmentation plugin.

### Transcript analysis

Snap frozen tumor pieces were placed into 1 mL TRIzol^TM^ reagent (Invitrogen) and mechanically dissociated. Following chloroform extraction and ethanol precipitation, RNA was quantified via NanoDrop. 1 μg of RNA was then converted into cDNA, and 10 ng/well cDNA was used for qPCR with TaqMan^®^ primer/probes for GAPDH, β2M, and IL-12β (ThermoFisher Scientific).

